# Auditory Feature-based Perceptual Distance

**DOI:** 10.1101/2024.02.28.582631

**Authors:** Shukai Chen, Marvin Thielk, Timothy Q. Gentner

## Abstract

Studies comparing acoustic signals often rely on pixel-wise differences between spectrograms, as in for example mean squared error (MSE). Pixel-wise errors are not representative of perceptual sensitivity, however, and such measures can be highly sensitive to small local signal changes that may be imperceptible. In computer vision, high-level visual features extracted with convolutional neural networks (CNN) can be used to calculate the fidelity of computer-generated images. Here, we propose the auditory perceptual distance (APD) metric based on acoustic features extracted with an unsupervised CNN and validated by perceptual behavior. Using complex vocal signals from songbirds, we trained a Siamese CNN on a self-supervised task using spectrograms rescaled to match the auditory frequency sensitivity of European starlings, Sturnus vulgaris. We define APD for any pair of sounds as the cosine distance between corresponding feature vectors extracted by the trained CNN. We show that APD is more robust to temporal and spectral translation than MSE, and captures the sigmoidal shape of typical behavioral psychometric functions over complex acoustic spaces. When fine-tuned using starlings’ behavioral judgments of naturalistic song syllables, the APD model yields even more accurate predictions of perceptual sensitivity, discrimination, and categorization on novel complex (high-dimensional) acoustic dimensions, including diverging decisions for identical stimuli following different training conditions. Thus, the APD model outperforms MSE in robustness and perceptual accuracy, and offers tunability to match experience-dependent perceptual biases.

## Introduction

Characterizing and comparing natural acoustic signals in a manner that mirrors perception is vital for researchers across wide-ranging fields spanning neuroscience, artificial intelligence, and psychology. These sounds, which include vocal and other acoustic communication signals as well as environmental sounds, typically vary simultaneously along multiple physical dimensions. Each of these dimensions may carry different (or no) behaviorally relevant information and which may vary across contexts. Species differences can complicate matters even further, as the features that carry perceptual relevance for one species may not necessarily generalize to another [1]. Thus, an ideal measure of the perceptually relevant similarities and differences between natural acoustic signals should be able to capture the complex multi-variate feature spaces of these sounds, and have a flexibility that permits tuning to context- and species-specific functional outputs.

To quantify differences between two audio signals, current studies, especially those using machine learning for audio generation, often resort to pixel-wise error functions between corresponding spectrograms, such as mean squared error (MSE) [2–4]. This approach has the benefit of easy quantification, but deviates from perception in significant ways. First, representations, spectrograms, commonly scale signal power across frequencies (Hz) linearly which is not representative of how frequency differences are perceived [5, 6]. In animals, including both humans and European starlings (Fig 1, frequency sensitivity, defined as the minimum detectable change in frequency, is not uniform across the frequency spectrum but scales positively with frequency [7]. That is, a 20 Hz frequency deviation at 8,000Hz is perceptually less noticeable than a 20Hz change at 1,000 Hz, although the absolute values of frequency deviations are the same. Log- or Mel-scaled spectrograms can mitigate this discrepancy somewhat, but fail to compensate fully for the non-uniformity of frequency sensitivity [8]. A second problem lies in the comparison of the signals. MSE and other pixel-wise error functions are naive to the statistical structure of the signals under comparison, and consider each time-frequency coefficient from the constituent spectrograms as an independent “feature”. As a result, these measures focus only on local (time-frequency restricted) details, weigh all such local deviations equally, and thus are highly sensitive to noise and other perceptually irrelevant signal perturbations. These instabilities are evident in Fig 1 where two spectrograms offset from each other by only one spectral slice, result in significant pixel-wise MSE across all frequency bands, whereas the perceptual difference between the two signals would be minimal. Finally, perceptual differences are not fixed. Vocal communication signals in particular are highly context-dependent. Human speech contains many examples where perception is biased strongly by language experience [9] or by local contextual cues within words, e.g. the Ganong effect [10]. A more ideal auditory distance metric would be more closely representative of perception, invariant to behaviorally irrelevant perturbation, and dynamically adjustable to context. Here we describe such a metric.

**Fig 1.**
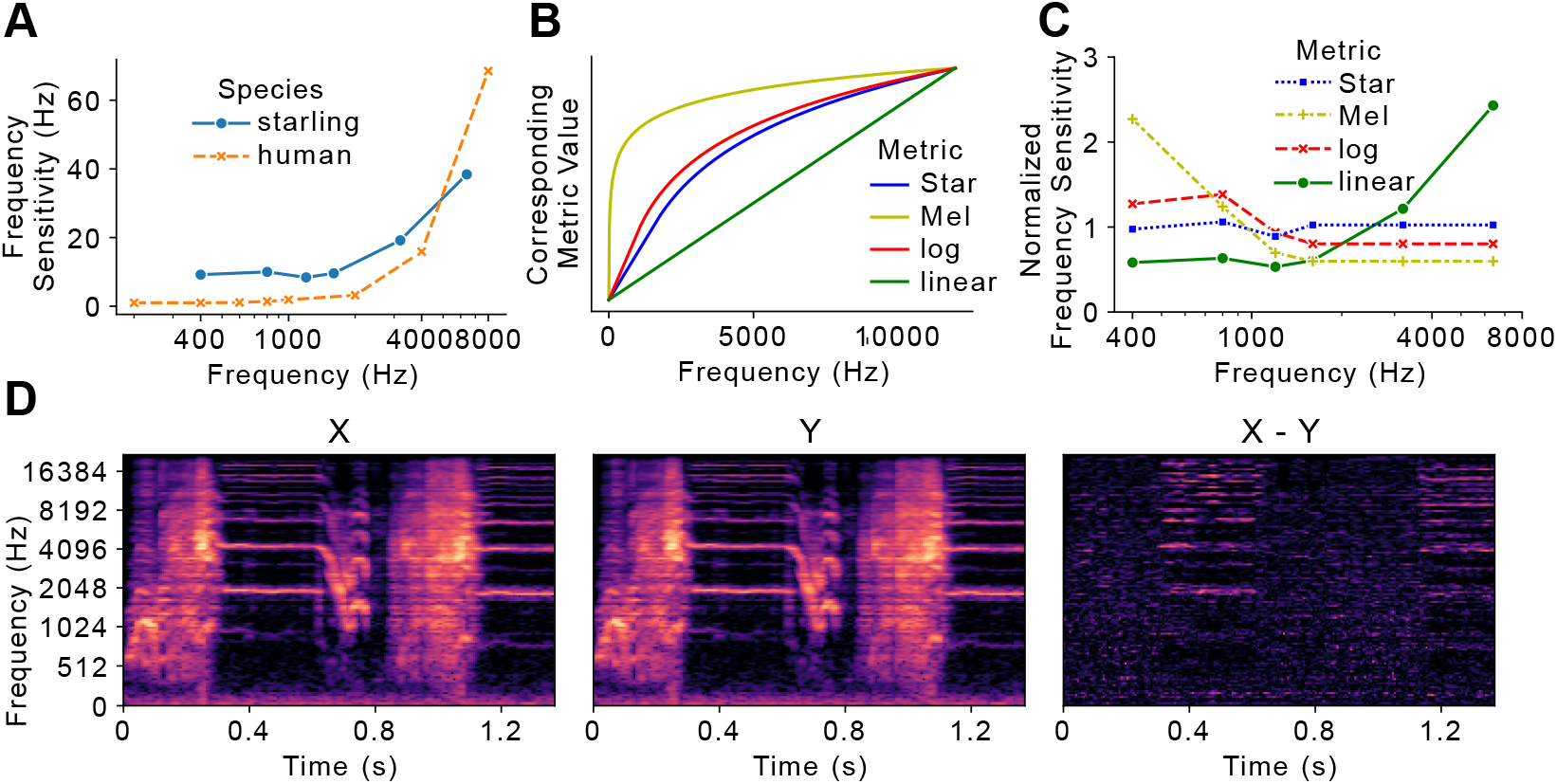
Current approach limitations. A) Frequency sensitivity, defined as the minimum discernable change at a given frequency, in both starlings and humans. B) Mapping between different frequency scaling metrics. C) Normalized frequency sensitivities in each frequency scale. Sensitivity values are normalized to the mean within each frequency scale. D) Two identical Star-scale spectrograms (X and Y) from each other by only one frequency band, corresponding to one Star. The distance spectrogram (X-Y) is calculated from pixel-wise subtraction between X and Y.

As an alternative to pixel-wise error, recent studies in computer vision quantify the visual difference (distance) between two images based on high-level features [11–14]. Instead of calculating per-pixel differences, these error functions extract and compare embeddings of images within the layers of convolutional neural networks (CNN). These embeddings successfully capture significant global features [11, 12], and can be used as quantifiable measures of perceptual distances between images [14]. For several deep neural networks, particularly those designed for image generation, feature vector based loss functions have enabled successful feature visualization [11], texture synthesis [15], and image style transfer [14]. Although powerful, the foregoing feature sets rely on highly labeled image datasets such as ImageNet [16], and are not likely to generalize across perceptual modality or task [17]. One solution to the constraint of hand-labeling is to apply self-supervised learning to unlabeled, readily available datasets [18–21]. Instead of using true labels as training targets, self-supervised training involves pretext tasks, which apply automatic preprocessing to an unlabeled dataset and optimizes on corresponding machine-generated pseudo-labels. A popular self-supervised algorithm for visual feature extraction is ”Jigsaw”, where an image is decomposed into small randomly-ordered puzzle pieces. When tasked to reorder shuffled pieces, the network eventually learns to identify significant visual features from images [19]. A similar algorithm has been proposed for spectrograms where the network learns acoustic features from reordering spectrogram fragments [22]. Unlike the original jigsaw model, the model on spectrograms performs the best when spectrograms are dissected in only the frequency domain.

Considering the forgoing challenges, constraints, and machine learning (ML) methods, we propose the Auditory Perceptual Distance (APD) metric, a computational method to quantify perceptual distances based on high-dimensional spectrotemporal features of acoustic communication signals. The APD metric combines innovative ML approaches to tackle the shortcomings of static pixel-wise error functions through a three-step process (Fig 2) optimized for auditory peripheral tuning accuracy, acoustic feature-based embedding, and experience-dependent perceptual bias. As a test case, we focus on the complex natural vocalization repertoire and auditory perceptual characteristics of European starlings, a species of songbird. We evaluate the absolute performance of the APD metric on behavioral data where ground truth perception is available, and compare its relative accuracy to MSE.

**Fig 2.**
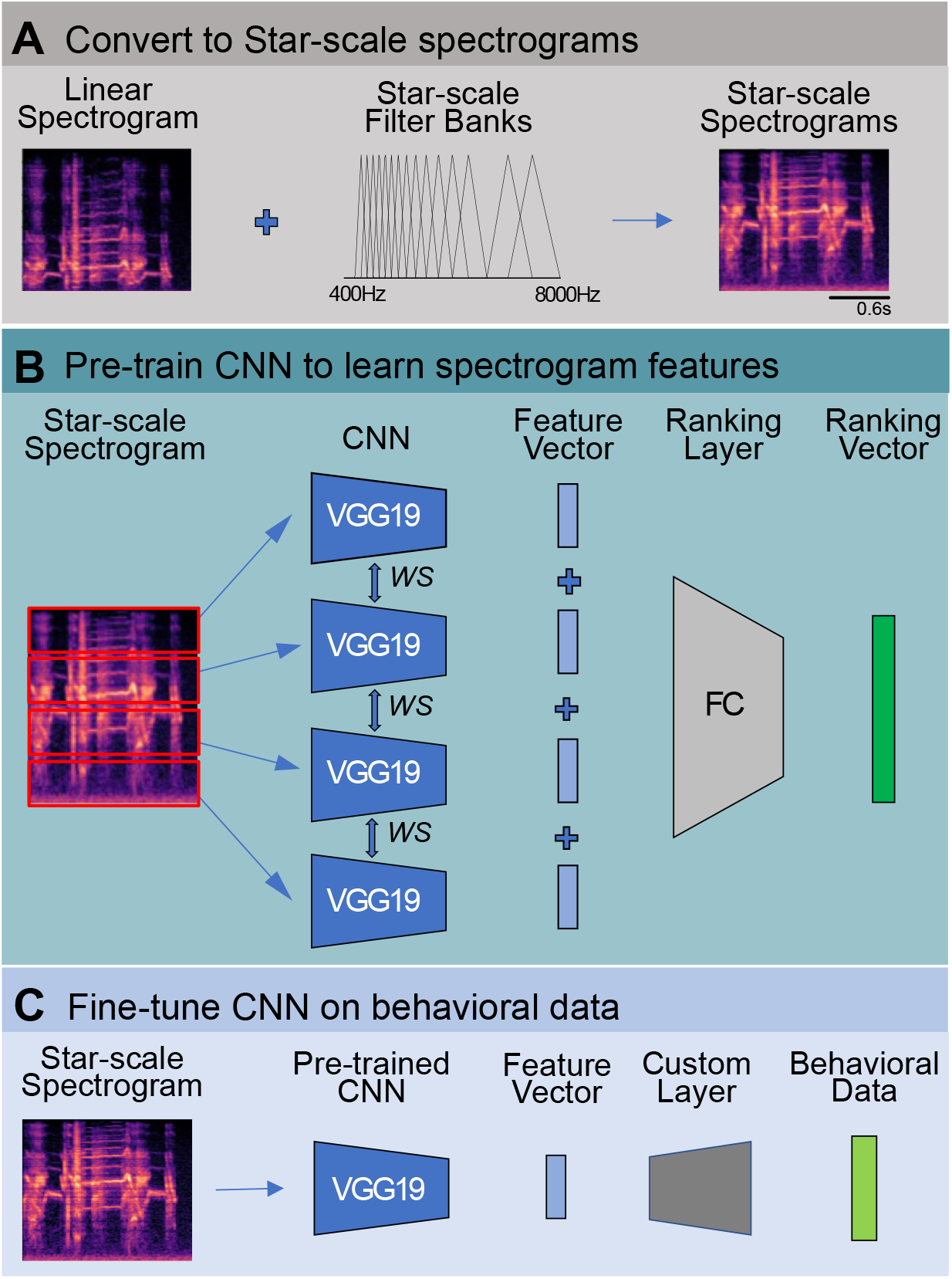
Three-step training process for the APD metric. A) Representation. Schematized process for converting to STAR-scale spectrograms. Linear spectrograms are converted to STAR-scale spectrograms through a set of STAR-scaled filter banks (see Materials and Methods for actual parameters). B) Pre-training. The CNN model is pre-trained on STAR-scale spectrograms of starling vocalizations to learn statistical features of song. Input spectrograms are divided into four equally sized spectral slices and shuffled (not shown here). The CNN outputs a ranking vector that indexes the original position of the shuffled slices. During training, all four slices are fed into the same CNN, yielding four feature vectors, which are then concatenated and passed to the fully connected (FC) ranking layers for classification. C) Fine-tuning. Once pre-trained, the learned CNN is fine-tuned on behavioral data. Pre-trained weights are transferred directly to the same CNN which is now connected to task-specific networks. Inputs to the fine-tuned model are unsegmented STAR-scale spectrograms. Outputs are animal judgments collected during the behavioral experiment.

## Results

Figure 2 outlines the three-step process for deriving the APD metric. Step one optimizes signal representation by devising the STAR-scale (Fig 2 A), a species-specific frequency scale, based on empirically measured critical bandwidths [7]. Mirroring similar scales in humans, such as the mel [5] and bark [23] scales, equal distances on the STAR-scale correspond to equal perceptual distance (Fig 1). We then train a CNN to learn significant acoustic features in a large library of starling songs using a self-supervised training algorithm (Fig 2 B, Materials and Methods). This process yields a naive embedding space that is familiar with vocal signals and the spectrotemporal features that characterize starling vocalizations statistically, but which lacks experience-dependent biases. To establish these biases, in step three we fine-tune the network-derived embedding space using behavioral data from real-world song perception experiments with starlings (Fig 2 C). In its final form, the APD metric enables characterization of vocal communication signals as low-dimensional feature vectors optimized for species and context-dependent perceptual experience. As a measure of the difference between signals in this perceptually-tuned embedding space, we take the cosine distance between acoustic feature vectors extracted by the trained CNN. We assess the effectiveness of each step separately and compare its performance to existing approaches. First, we inspect the perceptual uniformity of the STAR scale which we use to construct spectrograms for the APD metric. We then evaluate the effectiveness of pre-training on STAR-spectrograms compared to naive ImageNet weights, and the robustness of the pre-trained model to small local changes. Finally, we draw comparisons between computed and experimentally recorded perceptual distances after fine-tuning the metric on behavioral data, taking animal judgments as ground truth.

### STAR Scale

The APD metric is designed to work with spectrograms that capture species-specific frequency sensitivity. Ideally the frequency scale should be perceptually uniform. That is, any change of one unit should be perceived similarly by listeners regardless of absolute frequency values. To meet these requirements, we devise the STAR scale, a starling-specific frequency scale where equal distance on the STAR-scale represents equal perceptual distances observed (on average) by starlings (Fig 1). Accordingly, one “STAR” corresponds to the minimum detectable frequency difference at any given frequency. To compare the STAR scale to existing frequency scales, namely the Hz scale and the mel scale, we convert frequency sensitivity measurements collected by Kuhn *et al*. to units of STAR and mel (Fig 1) [7]. Sensitivity values are normalized to the mean within each frequency scale, as we focus more on the fluctuation of sensitivity across all frequency levels rather than the absolute values. Across the three frequency scales, the Hz scale shows the highest degree of variability (*µ* = 1.0, *σ* = 0.75, range 0.53∼2.43) followed by the mel scale (*µ* = 1.0, *σ* = 0.26, range 0.80∼1.38). The STAR scale is the most uniform (*µ* = 1.0, *σ* = 0.06, range 0.89∼1.06); even the maximum degree of fluctuation, logged at 120 STAR (1,200Hz), only measures 11%. Thus, the STAR scale outperforms both the Hz scale and the mel scale in terms of perceptual uniformity across frequencies.

### Pre-training

Estimating differences between spectrographic representations can be seen as an image processing problem. Following advances in computer vision, we compared embeddings of spectrograms within the layers of a well-known convolutional neural network (CNN), VGG-19 [24](Fig 2 C). Because spectrograms do not necessarily share the same behaviorally relevant feature space as images, we pre-train our network on STAR-scale spectrograms rather than adopting ImageNet weights like most computer vision models. Our pre-training dataset consists of 21,000 1.4s-long unlabeled starling vocalizations, converted to STAR-scale spectrograms (Fig 2, Materials and Methods). To guide learning we used ”Spectrographic Jigsaw” [19, 22], a spectrogram-specific adaptation of a popular self-supervised training task where networks learn high-level features under the pretext of sorting the shuffled puzzle pieces of an image (Fig 2). After the network is pre-trained in this way, the dense layers are disconnected from the CNN, the output of which is a 512-dimensional feature vector. fFinally, we randomly choose 30 1.4s long STAR-scale spectrograms for testing.

We evaluated the effectiveness of the STAR-scale spectrogram based APD metric. To distinguish this from subsequent versions that involve additional tuning to animal vocal judgements of vocalizations, we refer to the metric with only pre-training as the naive APD metric and the fine-tuned model (describe in the following section) as the tuned APD metric.

To assay robustness of the naive APD metric to shifts in time and frequency, we randomly chose 30 1.4s long STAR-scale spectrograms for testing. We consider two spectrograms offset by a variable number of rows or columns in either the frequency or the time domain, respectively. For simplicity, the offset was added as silence to one of the four edges of the spectrogram (top, bottom, left, right), and for each spectrogram, offset versions with opposite-edge offsets were compared (i.e., top vs. bottom, left vs. right). We expect the naive AP metric to react differently to temporal and spectral shifts. Namely, for two vocalizations offset only by trailing or leading silence similarity should be high, even when these offsets grow to many tens of milliseconds, as the relative differences int he spectro-temporal structure of the signal are unchanged. In contrast, while similarity for two vocalizations offset by only a few Hz should be high, the differences should increase as the frequency offset increases without saturating the metric. As shown in Fig 3, shifting by only a single (5ms) time step close to the gap detection threshold [25] for starlings yields a naive APD close to zero (APD=1.07E-4 ± 1.83E-4 normalized, N=30). Consistent with our intuition, the naive APD stays close to zero even with longer silence padding in the time domain (APD=5.93E-4 ± 8.65E-4 normalized, N=450). Shifts in the frequency domain (Fig.Fig 3) also conform to our intuition: a shift of one STAR, approximately the smallest frequency shift distinguishable by starlings, yields a very small shift in the naive APD (APD=3.16E-3 ± 1.46E-3 normalized, N=30) which steadily increases as the frequency offset grows. In contrast, MSE lacks a similar robustness to both temporal and spectral shifts. MSE is elevated significantly by even our smallest shift in time (5ms, MSE=0.06 ± 0.01 normalized, N=30) or frequency (MSE=0.06 ± 3.13E-3 normalized, N=30), and in both cases increases markedly for progressively larger shifts in each dimension.

**Fig 3.**
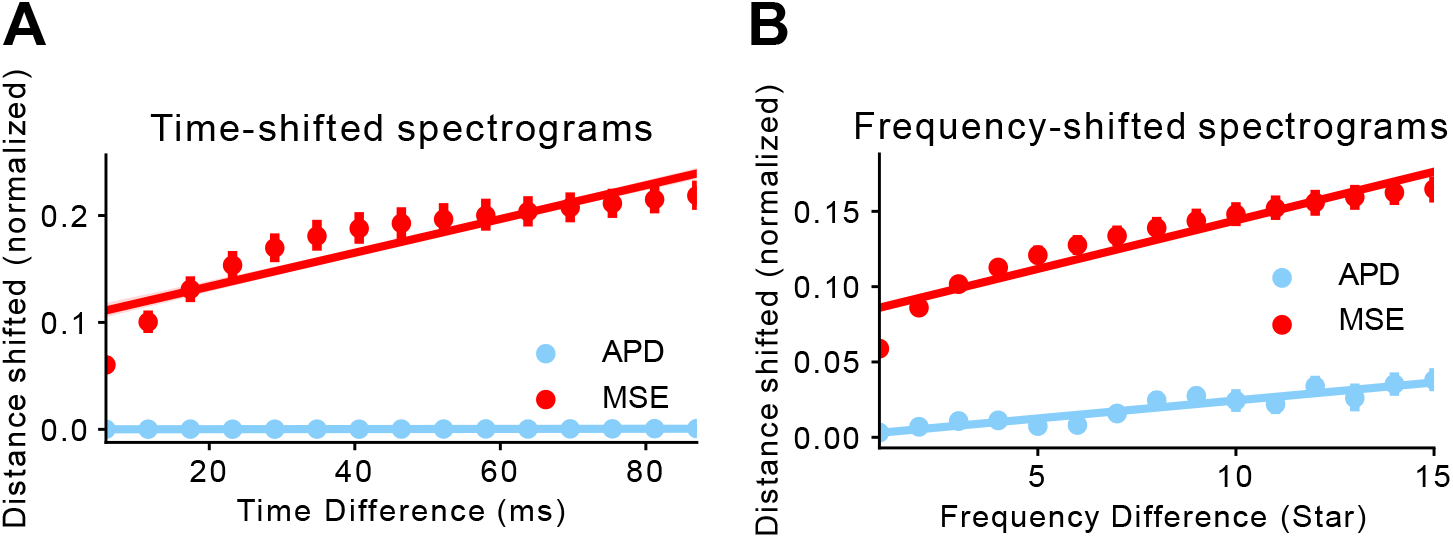
Perceptual loss is more robust to small changes than pixel-wise loss. We measure the distance between two time/frequency-shifted spectrograms using MSE and APD, and compare the trajectories of both distance metric as shifting becomes more significant.Both MSE and APD measurements have been normalized so that the maximum and minimum achievable distances are 1 and 0, respectively. A) Distance measured between two time-shifted spectrograms. We start with a pair of spectrograms separated by 5ms (leftmost points) and progress at a step size of approximately 5ms. We then calculate MSE and APD between each pair and fit a linear regression to all the points within either distance metric. B) Distance measured between two frequency-shifted spectrograms. Similarly, we start with a pair of spectrograms separated by 1 STAR (leftmost points) and progress at a step size of 1 STAR.

### Fine-tuning

The outcome of the pre-training process suggests that, compared to MSE, the naive APD model is better able to extract global features embedded in spectrograms consistent with intuition. The ultimate ground truth for modelling perceptual distance, however, is species-specific perceptual judgement. To better approximate real-world perception, we fine-tuned the naive APD metric on naturalistic artificial stimuli used in a behavioral experiment. The original experiment and its findings have been detailed elsewhere [26]. Briefly, starlings were trained on a two-alternative operant choice task to classify eight natural song syllables similar to those used to train the networks for naive APD; four syllables (e.g., A, B, C,and D) were associated with a one operant response and another four (E, F, G, and H) were associated with a second operant response. Once the subjects achieved stable recognition on these natural stimuli, a large set of artificial (“morph”) syllables were introduced using a double staircase procedure. Each morph syllable was a machine-generated linear interpolation between two of the natural syllables used for training (Fig 4, Materials and Methods). Subjects classified the artificial syllables based on their training with the natural syllables, yielding sixteen psychometric functions capture perceptual sensitivity on each of the 16 dimensions where syllables A-D were morphed to syllables E-H. Each subject was free to place the decision boundary between syllables (defined as the point of subjective equality) at any point along each syllable continuum.

**Fig 4.**
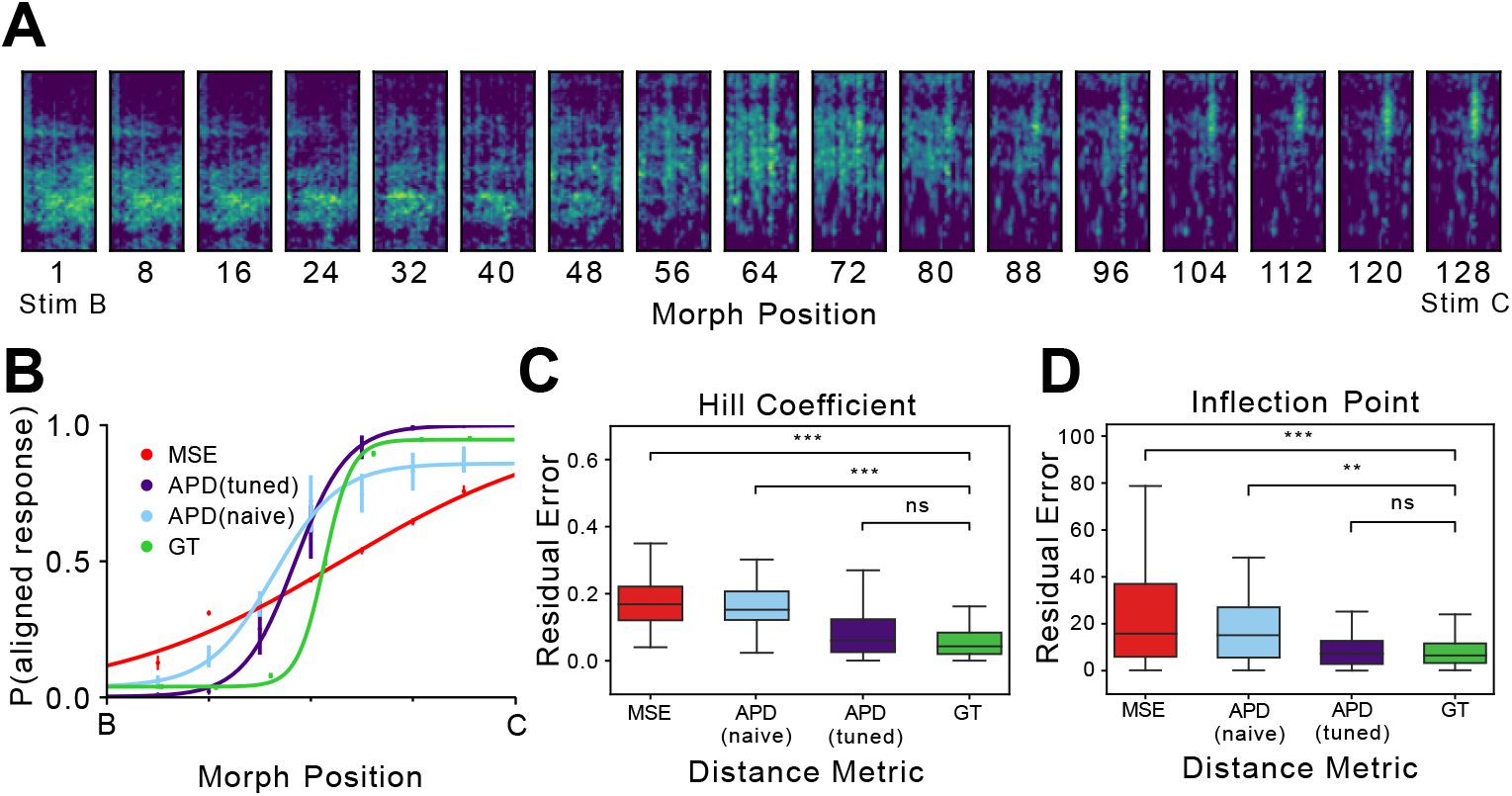
Fine-tuned APD outperforms both naive APD and MSE. A) An example set of morph stimuli, interpolated between stimuli B (left) and C (right). Refer to [26] for a detailed description of stimuli generation. Briefly, the author linearly interpolated between low-dimensional representations of B and C, and reverted the interpolation vectors to spectrograms. B) Computed and behaviorally measured psychometric curves on example morph stimuli shown in A. APD (naive) and APD (tuned) are both calculated from APD feature vectors, with the former only pre-trained (naive) and the latter fine-tuned on animal behavior data (tuned). C) Pairwise error in Hill coefficient measurements between each distance metric and the ground truth. For each computed psychometric curve (MSE, naive APD, and tuned APD), we calculate the error between its computed Hill coefficient and the ground truth value under the same training conditions (morph stimuli, cohort, etc.). For the ground truth, we calculate its internal variability by measuring errors between all pairs of subject judgments under the same training conditions. Outliers are not plotted. D) Pairwise error in inflection point measurements between distance metrics and ground truths. Error calculation follows the same pattern mentioned in C. All computed psychometric curves yield measurements within the variability of ground truths. Outliers are not plotted.

A double staircase training procedure was used to allow the birds to place their perceptual boundaries freely among all linear interpolated morph stimuli (Materials and Methods). One of the main findings was that birds trained under the same condition (for example, peck left for ABCD, right for EFGH) yielded a remarkable degree of consensus on decision boundaries across morph stimuli, suggesting a shared perceptual space. To replicate the training process computationally, we simulate a naive bird’s perceptual space with our naive APD model, and a trained bird’s with our fine-tuned APD model, trained on aforementioned experimental decisions made by birds. To assess the performance of the tuned APD model, we test our models on morph stimuli and draw comparisons between the APD-simulated and the experimentally measured psychometric curves.

Here we hypothesize that a high-performing perceptual distance metric achieves a high resemblance to the ground truth, specifically in terms of the inflection point and the Hill coefficient. The inflection point marks the decision boundary within a set of stimuli whereas the Hill coefficient entails sensitivity at the inflection point, measured as the slope of the psychometric curve [27–29]. In an example comparison between computed and measured psychometric curves (Fig 4), fine-tuned APD is capable of yielding simulations close to the ground truth whereas there exists an apparent mismatch between MSE and the ground truth, especially in sensitivity. A similar trend is observed across all stimuli sets for both computational distance metrics (Fig 4). To characterize resemblance to the ground truth, we calculate the absolute residual error between each computed Hill coefficient and the ground truth under the same training conditions (morph stimuli, cohort, etc.), and compare it to internal variability within the ground truth (GT), computed as absolute errors between all pairs of subject judgments under the same training conditions (Fig 4). The residual errors incurred by MSE are significantly different from the ground truth variability [*µ* = 0.160, *σ* = 0.058, compared to GT: p¡0.001, linear mixed effects model (LMM)] while the sensitivity of the fine-tuned APD curve is much closer to the ground truth [tuned APD: *µ* = 0.078, *σ* = 0.065, p¿0.1, LMM against GT]. The same set of residual errors is calculated for the inflection point (Fig 4). Similar results are observed: the tuned APD highly resembles the ground truth [tuned APD: *µ* = 8.76, *σ* = 7.85, p¿0.1, LMM against GT], whereas MSE shows significant deviation from the ground truth [*µ* = 16.17, *σ* = 13.62, p¡0.001, LMM against GT]. These results suggest APD is indeed a high-performing perceptual distance, yielding decision boundaries and sensitivities within the variability of the ground truth. Moreover, direct comparisons between APD and MSE provide a strong demonstration that APD is much more representative of starling perceptual sensitivity around the decision boundary than MSE.

Additional comparisons between naive and tuned APD models suggest that fine-tuning is imperative to achieving perceptually accurate simulations (Fig 4). Without fine-tuning, both sensitivity and decision boundary accuracy yielded by naive APD drop significantly lower than the ground truth [Hill coefficient: *µ* = 0.154, *σ* = 0.057, p¡0.01, LMM against GT; inflection point: *µ* = 15.48, *σ* = 11.97, p¡0.001, LMM against GT] whereas sensitivity level of the fine-tuned APD model is within the variability of the ground truth as mentioned earlier. These results indicate that accurate decision boundary placements and sensitivity characterization require fine-tuning on behavioral data.

Our preliminary results demonstrate that the APD model achieves high fidelity in characterizing starling perception. Through the characterization of frequency sensitivity at all frequency levels, we demonstrate that the STAR scale achieves higher perceptual uniformity than other existing frequency scales such as the Hz scale and the mel scale. By measuring the accumulation of error incurred through shifting spectrograms in both the spectral and the temporal direction, we find that APD successfully addresses the instability issue common in pixel-wise errors. And finally, by incorporating behavioral data into our training pipeline, we show that the tunable nature of APD significantly improves its ability to characterize animal perception. For each step of training the APD model, we systematically prove that the innovative approach involved directly leads to a better performance than the existing methods and therefore contributes to the observed high fidelity.

The significance of these preliminary findings is multifaceted. First, the introduction of the STAR scale answers the long-existing call for animal-specific perceptual scales. While starlings and humans share similar psychoacoustic abilities, using the Hz scale or the mel scale for research on starling perception is intrinsically problematic as neither is truly representative of starling’s unique peripheral frequency sensitivity.

The results also suggest that the incorporation of CNN to extract perceptual embedding is indispensable to the success of our model. Although there have been a few computer vision studies using CNNs to extract perceptual features, our model is the first to successfully apply this method to auditory perception. On one hand, the nature of a CNN predicate its success at extracting high-level features and consequently its robustness to local perturbations. Our results in experiment 2 indicate that the APD not only tackles MSE’s instability to small local changes but also offers different portrayals of the animal’s responses to temporal and spectral shifts. Given songbirds’ proven capability of processing relative pitch and relative timing [30, 31], such differential responses conform to our expectation: two audio signals differing only in the trailing or leading silence should contain resembling information whereas two signals separated by a few octaves should convey much more distinct messages. We argue the differential treatment originates from pre-training the CNN on spectrograms. Intuitively, the CNN learns to extract spectral features differently at various frequencies. In fact, if we directly use ImageNet weights, shifting in the frequency domain results in APDs close to zero, proving pre-training is essential for an accurate representation of songbirds’ ability to differentially respond to temporal and spectral shifts.

Compared to universal error metrics such as MSE, a CNN-based error metric offers the unparalleled advantage of tunability. Findings in experiment 3 exemplify the significant improvement in both sensitivity characterization and decision boundary placement only made possible by fine-tuning the APD model on animal judgments. In the original behavioral experiment, subjects were divided into cohorts where training conditions differ (Fig 5). Interestingly, a rare observation was that subjects in the same cohort arrive at similar decision boundaries on the same stimuli set while boundaries placed by different cohorts diverge (Fig 5) [26]. We show that through differential training to mimic different cohorts, the APD model can arrive at the same diverging boundaries as in the experimental data (Fig 5). While the naive APD fails to capture this property, its predictions between the two decision boundaries are scattered around 0.5, indicating uncertainty, whereas the predictions beyond both boundaries are much more determined, floating close to 0 or 1. This observation offers insight into what fine-tuning does to the model: we hypothesize that fine-tuning polarizes stimuli recognition by recalibrating response probabilities for both target categories to better align with the contrasting features between targets. In other words, fine-tuning realigns the originally global feature vector to reflect specific aspects of the stimuli and subsequently assigns a recognition threshold in that feature space. As a result, what is uncertain to a naive model can be tuned in either direction depending on the training condition, leading to the observed characterization of diverging decision boundaries. Intuitively, this process matches the effect behavioral training has on experiment subjects: while the subject already has an internal measure of perceptual distance before training, the training process teaches it to focus on specific features in the signal and iteratively refines the subject’s left/right decision thresholds based on feature distances. Another advantage of the APD model is its adaptability to the user’s task. Every step can be modified to fit the user’s specific need: the user has not only a wide range of pretext tasks for pre-training but also unrestrained freedom to fine-tune the model. The APD model can even be expanded to species beyond starlings. In the event of missing frequency sensitivity data to generate a species-specific frequency scale, the mel scale is an acceptable substitute, even showing similar results in starling perception thanks to its logarithmic nature. For starling perception, the STAR scale is still recommended as it is more perceptually accurate than the mel scale.

**Fig 5.**
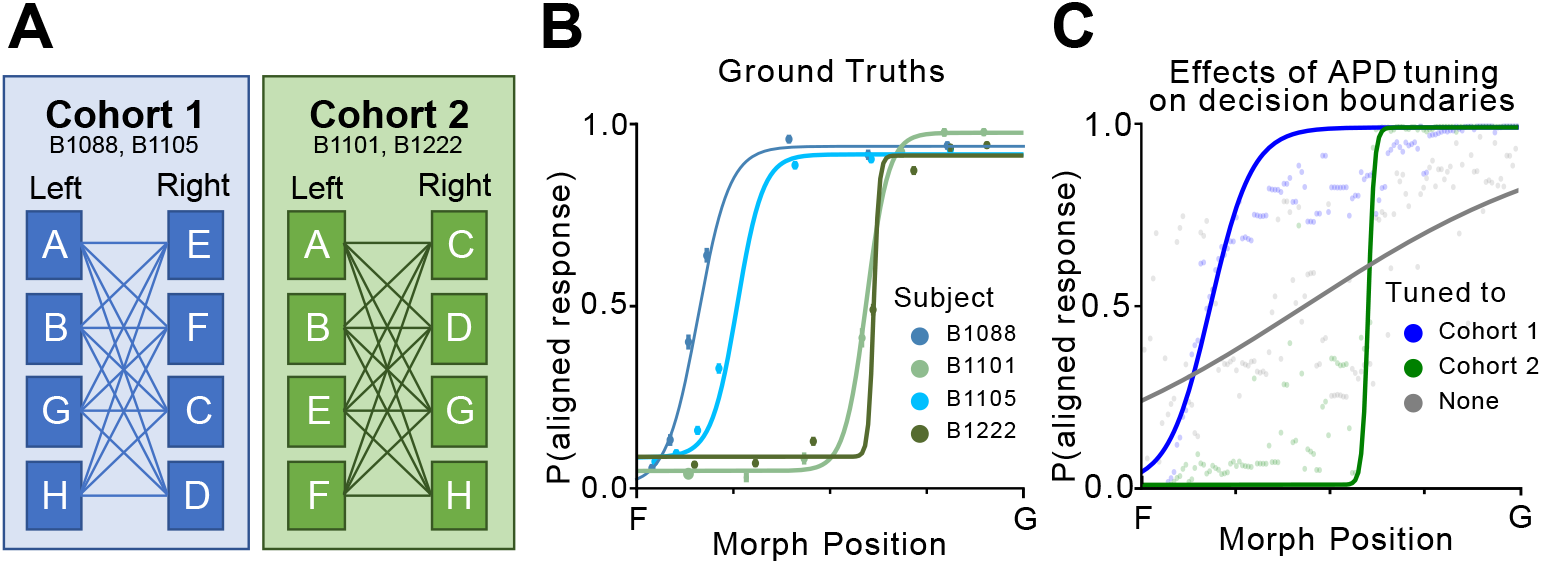
Fine-tuning realigns the model to a specific aspect of perception. A) Training schemes of two cohorts in the original experiment. Refer to [26] for a detailed description of the training process. Results from cohort 1 and 2 are shown with blue hues and green hues respectively in this figure. B) Behaviorally measured psychometric curves on morph stimuli between F and G. Each curve corresponds to a test subject in cohorts 1 and 2. C) APD-generated psychometric curves on morph stimuli between F and G. We compare results from APD models tuned on cohorts 1 and 2, as well as APD without fine-tuning (shown in gray).

Importantly, we believe that the APD model, while proven more perceptually accurate than existing methods, can be further optimized to achieve higher fidelity. A few components to tune include the CNN architecture, the pretext task employed during self-supervision, and the fine-tune training procedure. Our currently chosen approaches are direct adoptions from existing literature, but most hyperparameters have yet to be tuned for optimal performance because the tuning result would be highly specific to our data and less significant to other users looking to adopt the approach. Another future application is to use the APD model alongside a generative neural network for spectrograms as a loss function between the target and the generated spectrograms, similar to the loss function proposed by Johnson *et al*. [14]. Moreover, we acknowledge the possibility that the spectrotemporal information captured by the APD model from spectrograms, while already superior to MSE, could be further improved upon with other acoustic signal representations. One potential candidate is wavelet transforms, which have recently seen more usage in animal vocalizations [32–35]. Although currently we cannot say what the best solution is, we designed the APD model to be easily adaptable to any representation so it can be extensively studied and compared.

In this paper, we propose the APD model, a CNN-based model to quantify the perceptual distance between two auditory signals. Training the APD model is a three-step process, for each of which we systematically prove that the innovative approach involved directly leads to better performance than the existing methods and therefore contributes to better-portraying starling perception. Specifically, the APD model significantly outperforms MSE in terms of stability and sensitivity around perceptual boundaries and is therefore a more accurate representation of starling perception. In the future, we hope to optimize the APD model as well as use it as a loss function for generative neural networks for spectrograms.

## Materials and methods

### Datasets

#### Starling Vocalization Dataset

The dataset we use for pre-training the CNN was published by Sainburg *et al*. and available online [36]. It consists of songs from 14 European starlings individually collected in isolated chambers. All recordings were originally stored as 16 bit, 44.1 kHz wave files. From each singer’s hour-long recordings, we randomly segment 1,500 1.4s-long continuous vocalizations. The segmentation process is done automatically so that no syllable is truncated and no motif information is taken into consideration, meaning a signal can start and end inside a motif as long as there is no continuous silence longer than 0.5s.

#### Morph Stimuli Dataset

The morph dataset is directly borrowed from [26], where a more detailed description is available. Briefly, eight arbitrarily chosen motifs (labeled A H) are divided equally in three different ways (ABCD vs EFGH, ABGH vs EFCD, ABEF vs CDGH), forming a total of 24 unique pairs of motifs. Spectrograms of each pair of stimuli are passed through a trained autoencoder with a 64-dimensional bottleneck. Between the two 64-dimensional latent vectors, 128 linear interpolations are extracted, reverted back to full-size spectrograms using the same autoencoder, and subsequently inverted to wave files sampled at 48kHz using methods proposed by Griffin and Lim [37]. Altogether, the dataset consists of 3072 morph stimuli, including repeating endpoints.

### Behavioral Training Methods

#### Shaping

We adopt a multistage autoshaping routine [38] that familiarizes the birds with the apparatus, guides the bird to initiate trials, and associates trials with possible food rewards. On average, it takes the subjects 3-5 days to complete shaping, after which they start behavioral trials.

#### Baseline Training Procedure

All subjects learn to classify natural stimuli using a two-alternative choice (2AC) procedure [39]. Each subject initiates a trial by pecking at the center port on the panel, which triggers playback of a stimuli. The subject must peck the left or right port afterwards to indicate its choice. Each stimulus is associated with a ground truth, either left or right (4 each). Incorrect responses incur punishments (timeout), while correct responses result in rewards (food access). High response rates can be achieved, and stimulus-independent response biases can be ameliorated by manipulating the reinforcement schedules or introducing remedial trials according to established procedures. The subjects are trained with a variable reinforcement ratio of 4, meaning they need to get 1-7 (average of 4) correct choices in a row to be rewarded.

#### Double Staircase Procedure

Once the subject is able to classify natural stimuli at a high accuracy, we start the double staircase procedure to probe the perceptual boundary between each pair of left and right natural stimuli. The procedure works by estimating a window encompassing the boundary and iteratively reducing the window edge on either side based on the subject’s performance. The staircase procedure begins by randomly choosing one of the 16 possible natural stimuli pairs, and then selects a morph stimulus between the natural stimuli pair that is outside the window (90%) or just inside the window (10%). For an easy trial, the morph stimulus is one natural stimuli mixed only slightly with the other natural stimulus. For the probe trial, the stimulus is a morph just within the window the procedure believes the perceptual boundary to be in. If the subject gets a probe trial correct, the corresponding window edge advances to the location of the probe trial and further probe trials along this axis become more difficult. The subjects are rewarded by a variable reinforcement ratio mentioned above so that the birds are forced to perform on each trial but are not necessarily rewarded. This also allows for more trials per day.

### Computational Methods

#### STAR Scale

To convert a frequency value from the Hz scale to the STAR scale, the following formula is used:

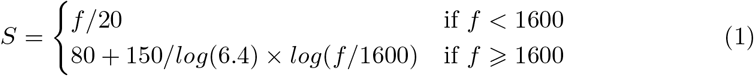

Where *f* is the frequency value in Hz. Note that the unit itself is meaningless, meaning it can be arbitrarily large or small depending on the multiplier attached to the formula.

#### Mel Scale

We use the mel scale conversion proposed by McFee *et al*. in Librosa [40].

#### Spectrogram Generation

All spectrograms used in this study are converted to the STAR scale. Computationally, this is a two-step process where Hz-scale spectrograms are first constructed from sound waves using an FFT size of 2048 combined with a step size of 256. Specifically for morph stimuli, spectrograms have a low-frequency cutoff of 850 Hz and a high-frequency cutoff of 10,000 Hz to avoid silence occupying the majority of the spectrogram. The Hz-scale spectrograms are then converted to STAR-scale spectrograms via a series of STAR filters, which are subsequently converted to decibel-based and normalized individually. All steps other than the STAR-scale conversion are accomplished with Librosa [40]. The final morph spectrograms are 186 STARs tall, whereas the final vocalization spectrograms are 291 STARs tall.

#### Model Architecture

Briefly, the APD model consists of a CNN and a few dense layers which are only used during training and dropped for feature extraction. Specifically for results included in this study, we use VGG19 as our choice of CNN due to the abundant literature on its capability of extracting high-level features [24]. During pre-training, we connect a single dense layer of size 4096 to the VGG19 whereas during fine-tuning, three dense layers (size 2048, 2048, 1024, respectively; ReLU activation) are used. Four separate input layers are connected simultaneously to the VGG19 during pre-training, rendering the model a Siamese network.

#### Pre-training

The APD model is pre-trained on a pretext task similar to the popular jigsaw task. For pre-training, we use the starling vocalization dataset in the form of STAR-scale spectrograms. 90% of the dataset is assigned the training set, whereas the remaining 10% is set aside as the validation set. There is no testing set because the purpose of pre-training is to transfer learned weights and thus we are uninterested in the pretext accuracy. Each spectrogram is divided in the frequency domain into four equally sized puzzle pieces which are subsequently shuffled. The goal of the model is to reorder the puzzle pieces based on relevant information extracted. During training, all four puzzle pieces are fed into the APD model, yielding four 512-dimensional vectors, which are then concatenated and passed to the dense layer for classification. The model is trained to minimize the MSE between target rank vectors and predicted rank vectors and is optimized with Adam (learning rate 10^*−*6^). A batch size of 32 is used in conjunction with a maximum of 1,000 epochs and early stopping to prevent overfitting, meaning the training will stop if validation accuracy does not improve.

#### APD Calculation

Once the model is trained, we can calculate APD between two spectrograms. This is achieved by first dropping the dense layers and reconfiguring the model so that there is only one input layer. Both spectrograms can be passed through the model, yielding two 512-dimensional feature vectors. We define the APD between these two spectrograms as the cosine distance between the two vectors, which ranges from -1 to 1, with a vaule of -1 indicating that the two vectors are exact opposites, 0 indicates they are orthogonal, and 1 indicates they are identical.

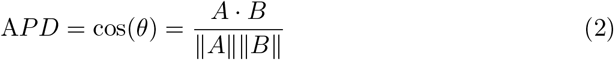

where *A* and *B* are CNN-derived feature vectors with *n* = 512 dimensions from the two spectrograms.

#### Shifting

To achieve the effect of shifting in either time or frequency domain, we pad the spectrograms with lines of silence. In the example of temporal shifting, we obtain two copies of the same spectrogram, one with n lines of silence added to the left and the other with n lines of silence added to the right. The APD model and MSE are then applied to both copies to characterize the error incurred by shifting the spectrogram.

#### Fine-tuning

We fine-tune the APD model on the morph stimuli dataset, under three different conditions to simulate the three cohorts in the behavioral experiment. For each set of morph stimuli judged by each subject, we train the model on all other morph stimuli sets using the subject’s response probability as target. For example, to investigate the model’s performance on mimicking subject X’s responses to stimuli set AE, we train a model on all remaining stimuli X has been exposed to, including AF, AG, AH, etc. If during behavioral experiment X classified *AF*_12_ as A 95% of the time, we assign a true label of [0.95, 0.05] to the stimulus. Starting with weights from the pre-training step, the model is optimized with Adam (learning rate 10^*−*6^) and trained to minimize categorical cross-entropy between true labels and predicted labels. To avoid overfitting, we inserted a 20% dropout after every dense layer. The output layer uses a softmax activation function to better characterize the structure of classification labels. Same as pre-training, early stopping is used to prevent overfitting.

#### Fitting Psychometric Curves

We model both the simulated and the measured behavior with a four-parameter logistic regression characterized by the following formula:

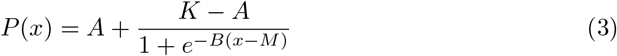

Where *A* and *K* are the minimum and the maximum value that can be obtained, respectively. *M* symbolizes the inflection point where the probability of yielding either response is 0.5. *B* represents Hill’s coefficient, the measured slope at the inflection point.

## Acknowledgements

Portions of this work were supported by NIH 5R01DC018055 to T.Q.G.

